# Pyruvate Oxidation Sustains B Cell Antigen-Specific Activation to Exacerbate MASH

**DOI:** 10.1101/2023.11.13.566832

**Authors:** Fanta Barrow, Haiguang Wang, Gavin Fredrickson, Kira Florczak, Erin Ciske, Shalil Khanal, Preethy Parthiban, Huy Nguyen, Enrique Rios, Enis Kostallari, Xavier S. Revelo

**Affiliations:** Department of Integrative Biology & Physiology, University of Minnesota Medical School, Minneapolis MN 55455, USA; Center for Immunology, University of Minnesota, Minneapolis MN 55455, USA; Division of Gastroenterology and Hepatology, Mayo Clinic, Rochester, MN 55905, USA

**Author notes:** Corresponding author. Xavier S. Revelo, Ph.D. Cancer & Cardiovascular Research Building 2231 6th St SE Minneapolis, MN 55455.

## Abstract

B cells play a crucial role in the pathogenesis of metabolic dysfunction-associated steatohepatitis (MASH), a severe form of steatotic liver disease that if persistent can lead to cirrhosis, liver failure, and cancer. Chronic inflammation and fibrosis are key features of MASH that determine disease progression and outcomes. Recent advances have revealed that pathogenic B cell-derived cytokines and antibodies promote the development of MASH. However, the mechanisms through which B cells promote fibrosis and the metabolic adaptations underlying their pathogenic responses remain unclear. Here, we report that a subset of mature B cells with heightened cytokine responses accumulate in the liver and promote inflammation in MASH. To meet the increased energetic demand of effector responses, B cells increase their ATP production via oxidative phosphorylation (OXPHOS) fueled by pyruvate oxidation in a B cell receptor (BCR)-specific manner. Blocking pyruvate oxidation completely abrogated the inflammatory capacity of MASH B cells. Accordingly, the restriction of the BCR led to MASH attenuation, including reductions in steatosis, hepatic inflammation, and fibrosis. Mechanistically, BCR restriction decreased B cell maturation, activation, and effector responses in the liver, accompanied by decreased T cell- and macrophage-mediated inflammation. Notably, attenuated liver fibrosis in BCR-restricted mice was associated with lower IgG production and decreased expression of Fc-gamma receptors on hepatic stellate cells. Together, these findings indicate a key role for B cell antigen-specific responses in promoting steatosis, inflammation, and fibrosis during MASH.

## INTRODUCTION

Metabolic dysfunction-associated steatotic liver disease (MASLD) is considered the liver manifestation of metabolic syndrome affecting approximately 32% of the adult population^1^. MASLD is a progressive disease that includes a wide spectrum of liver injury including simple steatosis and metabolic dysfunction-associated steatohepatitis (MASH) that may lead to serious complications such as liver cirrhosis and cancer. MASLD initiates with fat accumulation in the liver, which can progress to MASH, a more severe form of disease characterized by hepatocyte ballooning, inflammation, and varying degrees of fibrosis^2^. MASH has become the fastest-rising indication for liver transplantation^3^ and the leading cause of hepatocellular carcinoma (HCC)^4^. Despite these alarming rates, there are no FDA-approved therapies for the treatment of MASH. Inflammation plays a pivotal role in the progression of steatosis to the more severe MASH. Although the underlying mechanisms are poorly understood, several immune cells, including macrophages^5–8^, B cells^9–11^, CD4 T cells^12, 13^, CD8 T cells^14, 15^, neutrophils^16, 17^, and dendritic cells (DCs)^18^ have been shown to in promote inflammation and fibrosis during MASH.

Immune cell function is supported by metabolic adaptations that favor effector responses^19^. While naïve B cells require low energetic input to maintain quiescent homeostasis, activated B cells increase glucose uptake, glycolysis, and oxidative phosphorylation (OXPHOS) to meet the increased energy demand^20^. The microenvironment including the access to nutrients regulates the metabolic reprogramming in activated B cells, as observed in germinal center B cells that rely on fatty acid beta-oxidation for proliferation^21^. However, how the metabolic status of the MASH liver affects substrate utilization and energetic pathways in activated B cells, remains unexplored. Previously, we showed that pathogenic B cells accumulate in the MASH liver where they are activated by intestinal products via innate pattern recognition receptors in a process that was dependent on myeloid differentiation primary response 88 (MyD88) signaling^10^. However, B-cell activation during MASH also involved direct BCR stimulation, as shown using a reporter mouse in which BCR signaling induces green fluorescent protein expression under the control of the Nur77 gene^10^. While these data established the direct contribution of B cells to hepatic inflammation and injury in MASH, the mechanisms supporting B cell profibrogenic responses remained elusive. In the context of fibrosis, hepatic stellate cells (HSCs) are the major cell type responsible for fibrogenesis in the liver^22^. Upon activation, HSCs differentiate into collagen-producing myofibroblasts, resulting in collagen deposition and extracellular matrix remodeling^23^. Although we and others have shown that B cells have been shown to promote fibrosis in MASH^10, 11^, a critical gap exists in understanding whether B cells directly impact HSC activation and the specific mechanisms through which they contribute to liver fibrosis.

In this study, we show that B cells in the MASH liver have increased maturation and inflammatory cytokine production. To meet the energetic demand of heightened effector responses, MASH B cells selectively utilize pyruvate oxidation, and blocking this source of ATP abrogates their inflammatory potential in a BCR-dependent manner. Notably, BCR restriction protected against hepatic steatosis, inflammation, and fibrosis in association with lessened B cell-, T cell-, and macrophage-mediated hepatic inflammation. Thus, B cell BCR-specific responses are required for increased systemic and hepatic IgG antibodies, higher expression of Fc-gamma receptors on HSCs, and worsened fibrosis. Collectively, our findings demonstrate that MASH induces metabolic adaptations in B cells that favor antigen-specific responses that ultimately contribute to disease progression.

## RESULTS

### MASH induces a shift in intrahepatic B cell populations

To determine how MASH alters B cell populations in the liver, we fed C57BL6/J mice a high-fat high-carbohydrate (HFHC) diet for 15 weeks. We have previously shown that this regimen leads to the development of MASH, including a substantial increase in body weight, insulin resistance, hepatic steatosis, inflammation, and fibrosis^10^. Despite no changes in their proportion, MASH livers had an increased number of B cells, compared with controls (**Fig.1A**). Using RNAseq, we previously reported the presence of an immature B cell population characterized by a high expression of CD24^10^. Here, we used flow cytometry to determine the expression of CD24 in intrahepatic B cells from NCD and HFHC mice and found a decrease in CD24^hi^ B cells, consistent with our previous report^10^ (**Fig.1B**). However, high CD24 expression was not exclusive to immature B cells as other B cell populations also exhibited elevated CD24 levels **(Extended data** **Fig.1A**). To further validate the presence of immature B cells in the liver, we utilized IgM and IgD expression to characterize different B cell subsets based on their B cell receptor. This approach identified a decrease in the proportion of immature B cells in MASH, while mature B cells increased (**Fig.1C**). These changes were not observed in the spleen, suggesting tissue-specific effects likely driven by the liver microenvironment **(Extended data** **Fig.1B**). There were no alterations in the proportion of naïve, mature activated, and class-switched B cells. However, the number of class-switched and naïve B cells in the MASH liver was increased, compared with controls (**Fig.1C**). Next, we assessed how MASH affects antibody-producing B cells, previously shown to participate in the pathogenesis of MASH^9, 24^. There were no alterations in the number of plasmablasts and plasma cells within the MASH liver (**Fig.1D**). Taken together, our results indicate that the MASH liver contains distinct B cell subsets with a subset of B cells showing increased maturation during MASH.

**Figure 1.**
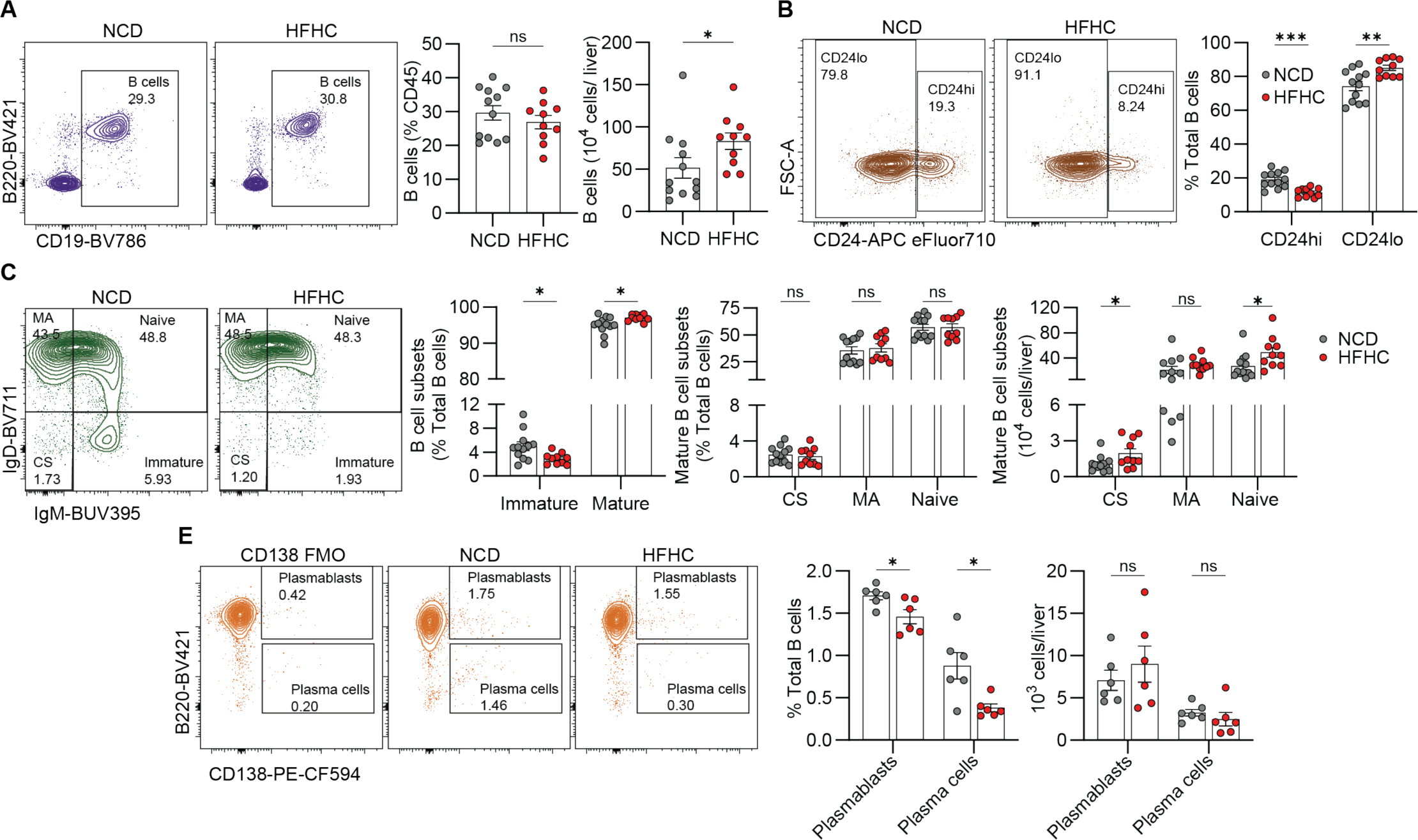
MASH induces a shift in intrahepatic B cell populations. **(A-E)** C57BL/6J mice were fed an NCD or MASH-inducing HFHC diet for 15-19 weeks. **(A)** Representative flow cytometry plot showing intrahepatic B cells (left), frequency (middle) and number (right). **(B)** Representative flow cytometry plot showing CD24^high^ and CD24^low^ B cell populations in the liver (left), and frequency (right). **(C)** Representative flow cytometry plot showing class-switched (CS), mature activated (MA), naïve, and immature B cells (far left) and frequency of total mature (CS, MA, and naïve) and immature B cell subsets (middle left), followed by frequency (middle right) and number (far right) of mature B cell subsets. **(D)** Representative flow cytometry plot showing antibody-secreting B cell subsets, plasma cells, and plasmablasts (left panels)followed by their frequency (middle) and number (right) in the liver. Data are representative of two independent experiments, ns = not significant, *p ≤ 0.05; **p ≤ 0.01; ***p ≤ 0.001; ****p ≤ 0.0001, unpaired t-test. Abbreviations: NCD, normal chow diet; HFHC, high-fat high-carbohydrate; CS, class-switched; MA, mature activated.

To investigate how MASH alters the production of antibodies by B cells, we purified intrahepatic B cells from NCD and HFHC mice and determine their antibody secretion in response to LPS or IgM/CD40 treatment to mimic toll-like receptor 4 (TLR4) and BCR stimulation, respectively. B cells from HFHC mice produced fewer antibodies in response to LPS (**Fig.2A**) and IgM/CD40 (**Fig.2B**). While we did not observe differences in the amount of IgA in the liver, we noted an increase in the amount of total IgG in the HFHC liver, compared with controls (**Fig.2C**). Similarly, we found increased IgG2a, IgG2b, and IgA in the serum of MASH mice (**Fig.2D**). Next, to characterize B cell cytokine responses, we employed Isoplexis, a single-cell-based secretome analysis assay. We isolated and purified B cells from NCD or HFHC livers, primed the cells with LPS for 16 hours, and analyzed their cytokine profiles. B cells from MASH mice exhibited a distinct cytokine profile characterized by a high polyfunctional strength index, which represents their ability to produce multiple types of cytokines **(Fig.2E, 2F)**. This assay showed that B cells from HFHC mice produced increased chemoattractive, effector, stimulatory, and inflammatory cytokines while B cells from NCD mice produced fewer and regulatory cytokines that dampen inflammation (**Fig.2F**). The primary cytokines produced by B cells from HFHC mice were RANTES, IL-6, IL-7, and Granzyme B (**Fig.2G**). In summary, the MASH liver accumulates IgG antibodies and harbors B cells with increased cytokine effector responses, while the production of antibodies induced by LPS and IgM/CD40 appears to be attenuated.

**Figure 2:**
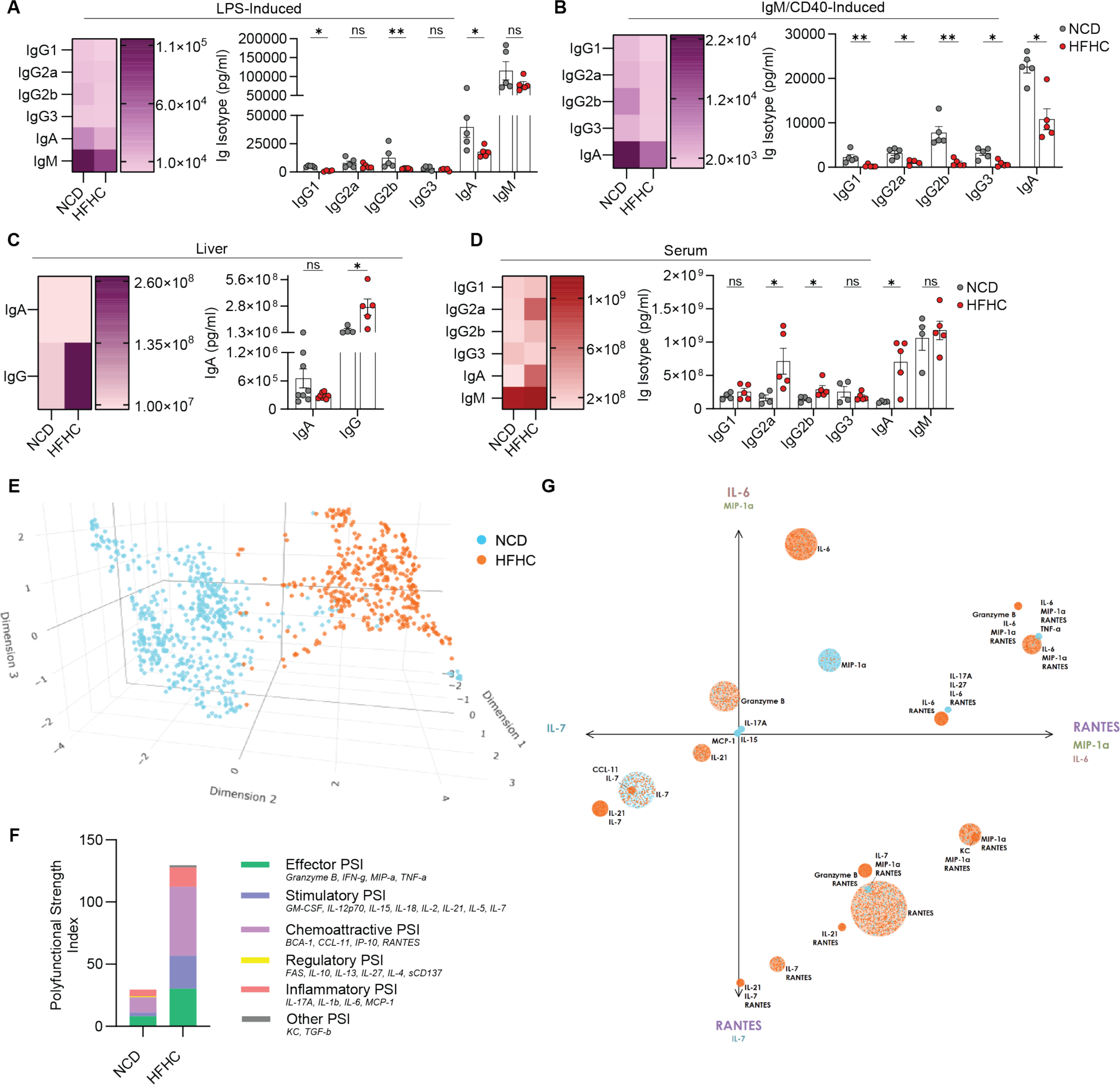
Increased IgG deposition in the liver during MASH. **(A-G)** Data is from C57BL/6J mice fed an NCD or HFHC diet for 15-20 weeks. **(A)** Heat map and quantification of immunoglobulin isotypes produced by purified intrahepatic B cells stimulated with LPS for 48 hours. **(B)** Heat map and quantification of immunoglobulin isotypes produced by purified intrahepatic B cells stimulated with IgM/CD40 for 48 hours. **(C)**. Heat map and quantification of IgA and IgG levels in liver tissue. **(D)** Heat map and quantification of immunoglobulin isotypes in the sera of NCD and HFHC mice. **(E-F)** Intrahepatic B cells were primed with LPS for 16 hours and then subjected to single-cell secretome analysis. **(E)** 3D t-SNE plot showing key differences in cytokine profiles between NCD and HFHC B cells. Each dot corresponds to a single cell. **(F)** Stacked bar graph showing the PSI, calculated by combining the polyfunctionality of the sample and the average intensity of the cytokines secreted by polyfunctional single cells. The height of each bar corresponds to the PSI of the corresponding group. The bars are broken down by the PSI of each cytokine class. **(G)** PAT PCA plot showing the functional and polyfunctional NCD and HFHC B cells and their combinatorial cytokine secretions. Dots represent single cells, and broader circles are color-weighted to the color of the group that secreted that group of cytokines with the highest frequency. The axes correspond to the principal components of the data, linear combinations of the specified cytokines; the cytokines most strongly present in each component are listed at the ends of the axes. Data are representative of two independent experiments except **E-F**, ns = not significant, *p ≤ 0.05; **p ≤ 0.01; ***p ≤ 0.001; ****p ≤ 0.0001, unpaired t-test. Abbreviations: NCD, normal chow diet; HFHC, high-fat high-carbohydrate; PSI, polyfunctional strength index; PAT, Polyfunctional Activation Topology.

### Intrahepatic B cells show increased ATP production during MASH

Given the heightened maturation and cytokine responses of B cells in MASH, we aimed to characterize the metabolic profile of B cells in the MASH liver and its connection to B cell responses by assessing key cellular respiration parameters. Using a real-time ATP rate Seahorse assay, we found that MASH B cells exhibit increased ATP production, primarily driven by the OXPHOS and to a lesser extent glycolysis (**Fig. 3A**). In support of this finding, we also found that MASH B cells display an elevated mitochondrial mass (**Fig. 3B**). To further confirm that MASH B cells depend on the OXPHOS pathway, we conducted a Mito stress test to assess the capacity of B cells to meet energetic demands. MASH B cells exhibit higher basal, ATP-linked, maximal respiration, and spare respiratory capacity oxygen consumption rates, as a result of their greater ability to meet energetic demands (**Fig. 3C**). Next, we explored which substrates contribute to the increased ATP production in MASH B cells. Using inhibitors targeting fatty acid oxidation (FAO), glutamine oxidation, and glucose and pyruvate oxidation, we examined the reliance of MASH B cells on these fuels to meet their higher energy requirements. Given the abundance of fatty acids in the MASH liver, we initially hypothesized that MASH B cells would rely on fatty acid oxidation. However, Etomoxir, an inhibitor of FAO that blocks carnitine palmitoyl-transferase 1a (CPT1a), did not reduce basal oxygen consumption rates or the capacity of the B cells to meet energetic demands under conditions of maximal respiration (**Fig. 3D**). As Etomoxir is specific for long-chain fatty acid oxidation, we also investigated whether short-chain fatty acids from the oxidation of very long-chain fatty acids in the peroxisome contributed to the increase in ATP production by MASH B cells. However, Thioridazine, a peroxisomal fatty acid oxidation inhibitor, had no effect on the basal oxygen consumption rate and spare respiratory capacity of MASH B cells **(Extended Data** Fig. 2A). Furthermore, we assessed the contribution of exogenous fatty acids to the elevated ATP production of MASH B cells with the use of FAO Blue, a fatty acid analog coumarin dye that enables the visualization of FAO^25^. This assay showed that MASH B cells engage in minimal fatty acid oxidation compared to healthy B cells, confirming that FAO is not a major oxidative pathway for MASH B cells **(Extended Data** Fig. 2B). We also explored the reliance of MASH B cells on glutamine oxidation to meet their heightened energy demand and found that inhibition of glutaminase 1 using BPTES revealed no difference in basal oxygen consumption rates or spare respiratory capacity for MASH B cells (**Fig. 3E**). Thus, we examined the role of pyruvate oxidation using UK5099, an inhibitor of pyruvate transport into the mitochondria for the TCA and OXPHOS pathways. We observed no difference in basal oxygen consumption rates but a significant attenuation in the ability of MASH B cells to meet energetic demands under maximal respiration conditions (**Fig. 3F**). These data indicate that MASH B cells selectively rely on pyruvate oxidation to boost their ATP production.

**Figure 3:**
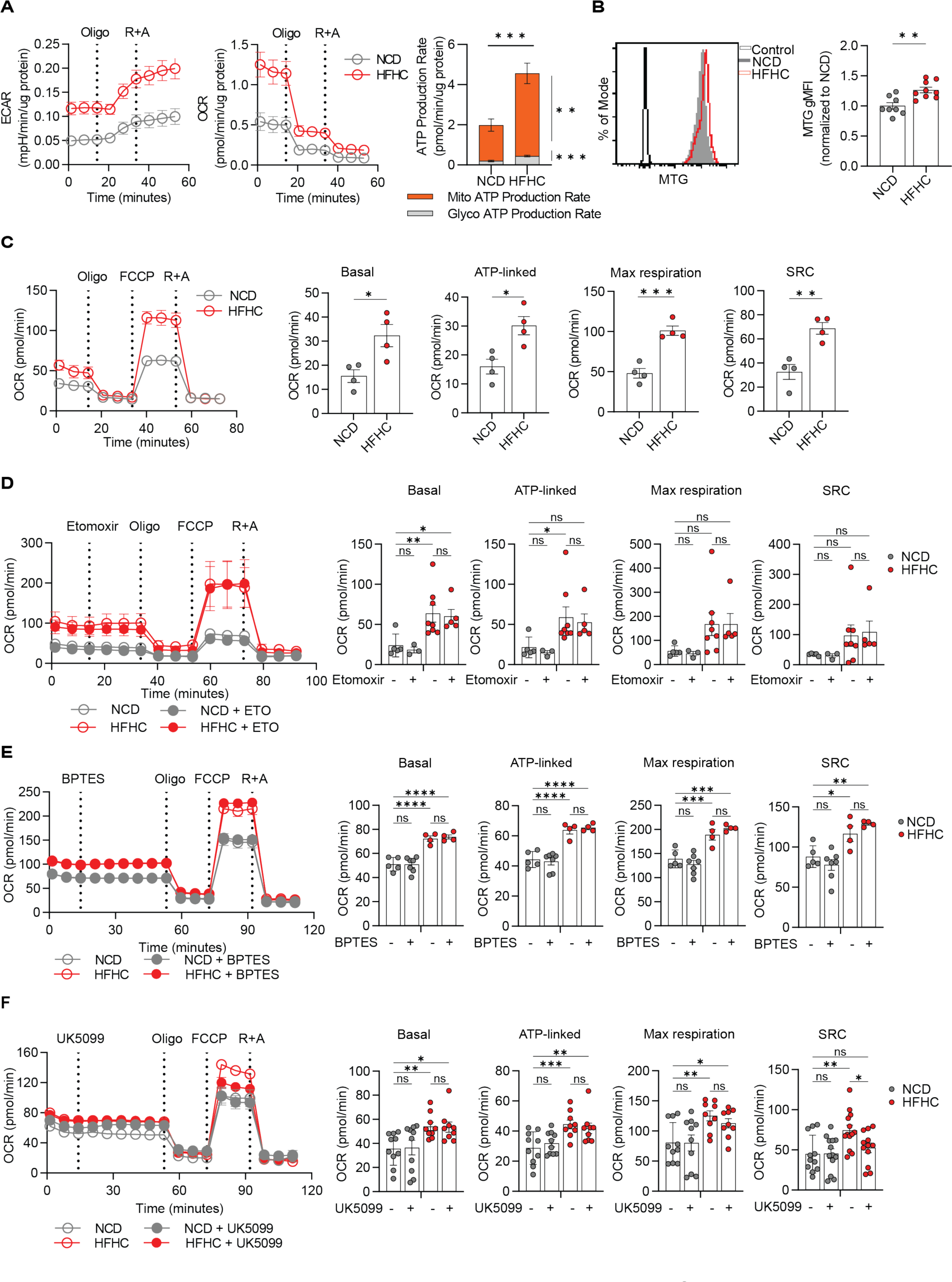
Intrahepatic B cells show increased ATP production during MASH. **(A-F)** Data is from C57BL/6J mice fed an NCD or HFHC diet for 15-21 weeks. Intrahepatic B cells were purified and subjected to extracellular flux analysis using the Seahorse XFe96 flux analyzer. **(A)** Representative trace of extracellular acidification rate (ECAR) (left), and oxygen consumption rate (OCR) (middle) followed by mean ATP production rate (right) of NCD and HFHC B cells. **(B)** Flow cytometry analysis of mitochondrial mass using MitoTracker Green (MTG), representative MFI plot (left), and quantification (right). **(C)** Representative trace and quantification of intrahepatic B cells subjected to a Mito stress test to gauge basal respiration, OCR linked to ATP production, maximal respiration, and spare respiratory capacity (SRC). SRC is the difference between basal OCR values and maximal OCR values obtained after FCCP uncoupling. **(D)** Representative trace and quantification of basal, ATP-linked, maximal, and SRC of intrahepatic B cells subjected to a substrate oxidation stress test with fatty acid oxidation inhibition using Etomoxir. **(E)** Representative trace and quantification of basal, ATP-linked, maximal, and SRC of intrahepatic B cells subjected to a Glutamine oxidation stress test with the glutaminase inhibitor BPTES. **(F)** Representative trace and quantification of basal, ATP-linked, maximal, and SRC of intrahepatic B cells subjected to a substrate oxidation stress test where pyruvate oxidation is inhibited using the mitochondrial pyruvate carrier inhibitor UK5099. Results represent the mean of 2 independent experiments. Bars represent mean +/− SEM; ns = not significant; *p ≤ 0.05; **p ≤ 0.01; ***p ≤ 0.001; ****p ≤ 0.0001 by One-Way ANOVA. Abbreviations: NCD, normal chow diet; HFHC, high-fat high-carbohydrate; Oligo, oligomycin; R+A, rotenone and antimycin A; SRC, spare respiratory capacity; FCCP, Carbonyl cyanide-4 (trifluoromethoxy) phenylhydrazone; SRC, spare respiratory capacity; MFI, mean fluorescence intensity.

### Pyruvate oxidation selectively regulates B cell effector responses during MASH

To explore the relationship between pyruvate metabolism and B cell function, we investigated the role of pyruvate oxidation in regulating B cell effector responses during MASH. We isolated B cells from MASH or healthy livers and stimulated them with lipopolysaccharide (LPS) or IgM/CD40. We found no differences in the production of IL-6 and TNFα by MASH or healthy B cells when treated with LPS and the UK5099 inhibitor **(Fig. 4A, 4B)**. Moreover, UK5099 did not impact LPS-induced antibody production by MASH B cells but it reduced IgG1 and IgM antibody production in B cells from NCD mice (**Fig. 4C**). Notably, when MASH B cells were stimulated with IgM/CD40 and treated with UK5099, there was a complete suppression of IL-6 and TNFα production in HFHC B cells **(Fig. 4D, 4E)**. IgM-CD40-induced IgG2a and IgA production by MASH B cells was not significantly affected despite trending decreases after treatment with the inhibitor (**Fig. 4F**). Collectively, these findings demonstrate that MASH B cells selectively rely on pyruvate oxidation to fuel their energetic demand of activation and effector responses in a BCR signaling-dependent manner.

**Figure 4:**
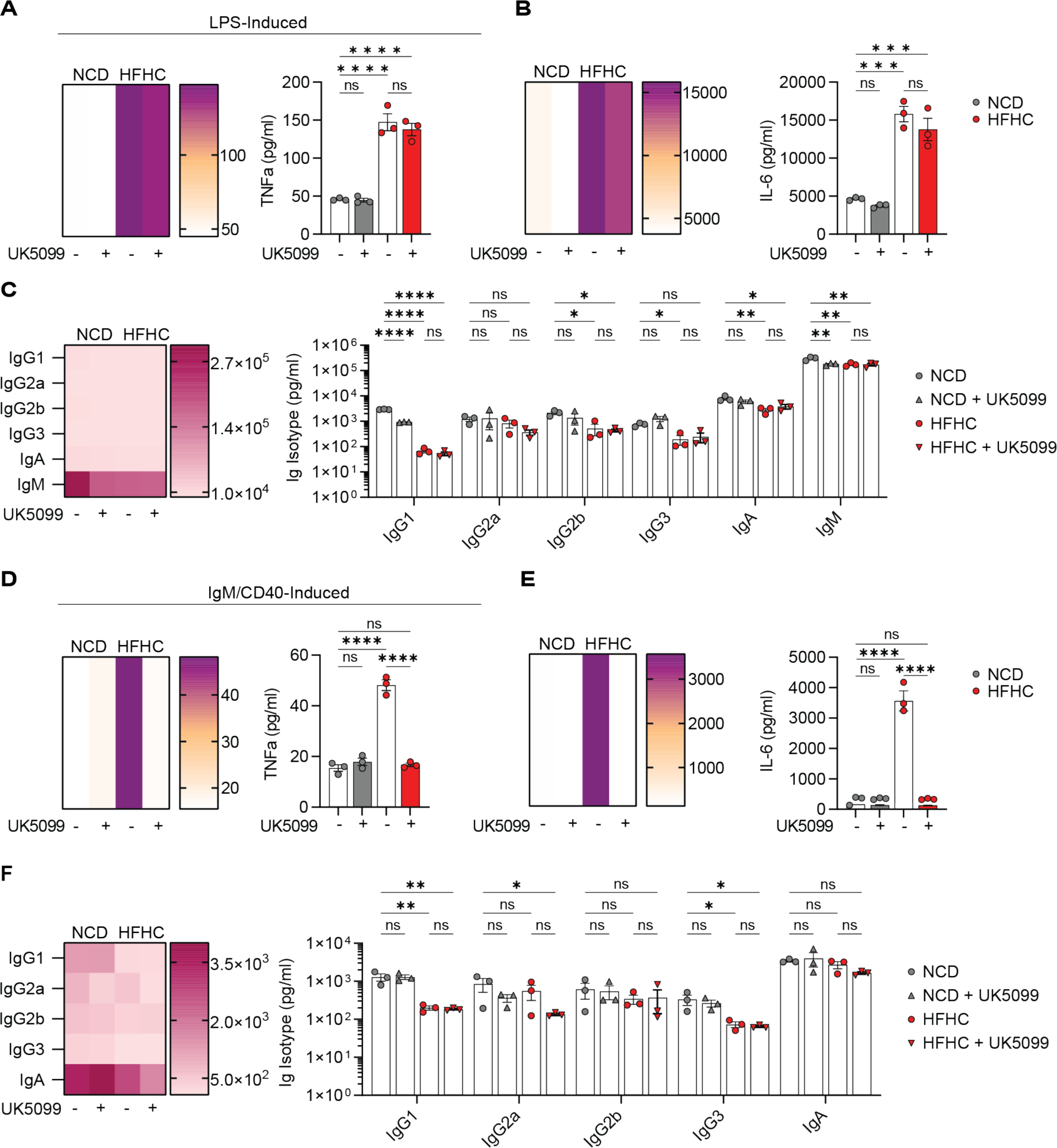
Pyruvate oxidation selectively regulates B cell effector responses during MASH. **(A-F)** Data is from C57BL/6J mice fed an NCD or HFHC diet for 15-25 weeks. Intrahepatic B cells were isolated from NCD or HFHC livers and treated with LPS or IgM/CD40 for 48 hours in the presence (triangle) or absence (circle) of the pyruvate oxidation inhibitor UK5099. Data is presented as heat maps and quantification of mean data. **(A)** LPS-induced TNF-α production. **(B)** LPS-induced IL-6 production. **(C)**. Immunoglobulin isotypes produced by LPS-stimulated intrahepatic B cells. **(D)** IgM/CD40-induced TNF-α production. **(E)** IgM/CD40-induced IL-6 production. **(F)**. Immunoglobulin isotypes produced by IgM/CD40-stimulated intrahepatic B cells. Data are representative of one independent experiments (n=16 NCD mice, n=8 HFHC mice; samples pooled to n=3 per group), ns = not significant, *p ≤ 0.05; **p ≤ 0.01; ***p ≤ 0.001; ****p ≤ 0.0001, One-Way ANOVA. Abbreviations: NCD, normal chow diet; HFHC, high-fat high-carbohydrate; LPS, lipopolysaccharide.

### B cell receptor restriction ameliorates MASH progression

Having established that MASH intrahepatic B cells rely on pyruvate oxidation to support their BCR-stimulated effector responses, we sought to investigate the role of antigen-specific responses via the BCR. We fed mice with a transgenic BCR restricted to recognize the irrelevant antigen hen egg lysozyme (HEL) not present in the mouse^26^ either a NCD or HFHC and determine their disease progression. HFHC-fed HEL mice showed no differences in body weight compared to controls (**Fig. 5A**) but had decreased liver weight (**Fig. 5B**), triglyceride content (**Fig. 5C**), and steatosis (**Fig. 5D**). Correspondingly, HEL mice had lower levels of ALT and AST, enzymes released by hepatocytes during liver injury **(Fig. 5E, 5F)**. Improvement in steatosis was accompanied by increased expression of genes involved in fatty acid oxidation in HEL livers compared to wild-type controls (**Fig. 5G**). No differences were observed in the expression of genes involved in lipogenesis between HEL and wild-type livers **(Extended Data** Fig. 3A), suggesting no perturbations in de novo lipogenesis. Next, we characterized the immune composition of HEL and wild-type livers using cytometry by time of flight (CyTOF) and flow cytometry. No differences were found in the proportion and number of B cells, CD4 T cells, CD8 T cells, natural killer (NK) cells, DCs, and neutrophils between HEL and wild-type livers **(Fig. 5H, Extended data** **Fig. 3B**). However, HEL intrahepatic B cells displayed an increased proportion of immature and naïve subsets accompanied by substantial decreases in mature activated and class-switched subsets (**Fig. 5I**). B cells with BCR restriction showed a decrease in their expression of IL-6 controls upon phorbol-myristate-acetate (PMA) stimulation (**Fig. 5J**). As B cell crosstalk with T cells has been shown to promote MASH^10, 14, 27^, we investigated the changes in T cell subsets in HFHC-fed HEL and wild-type control mice. The liver of HEL mice had CD4 T cells with fewer effector memory subsets (**Fig. 5K**), suggesting a decrease in activation. We also observed a decrease in the proportion of T helper 1 and regulatory T cells **(Extended Data** Fig. 3C), and a decrease in interferon-gamma (IFN-γ)^+^ and TNF-α^+^ CD4 T cells in the livers of HEL mice. Similarly, CD8 T cells had a lower frequency of naïve and central memory subsets **(Extended Data** Fig. 3D) and decreased in IFN-γ^+^ and TNF-α^+^ CD8 T cells (**Extended Data** Fig. 3E). We also investigated if inhibition of B cell antigen-specific responses affects macrophages during MASH. HEL livers had a higher proportion and number of embryonic-derived Kupffer cells (emKCs) and monocyte-derived VSIG4^-^ Tim4^-^ non-KCs, while no differences in moKCs and monocyte-derived macrophages (MoMFs) were observed (**Fig. 5M**). All macrophage subsets had decreased expression of CD80 (**Fig. 5N**), while emKCs, moKCs, and nonKCs had less CD86 expression (**Fig. 5O**), and emKCs and MoMFs had decreased major histocompatibility complex II (MHC-II) expression (**Fig. 5P**) indicating that the macrophage subsets in the HEL liver decrease their antigen presentation machinery compared to controls. Collectively, these findings demonstrate that BCR-mediated B cell activation promotes MASH directly by driving inflammation and indirectly by increasing steatosis and T cell and macrophage activation and inflammatory responses.

**Figure 5:**
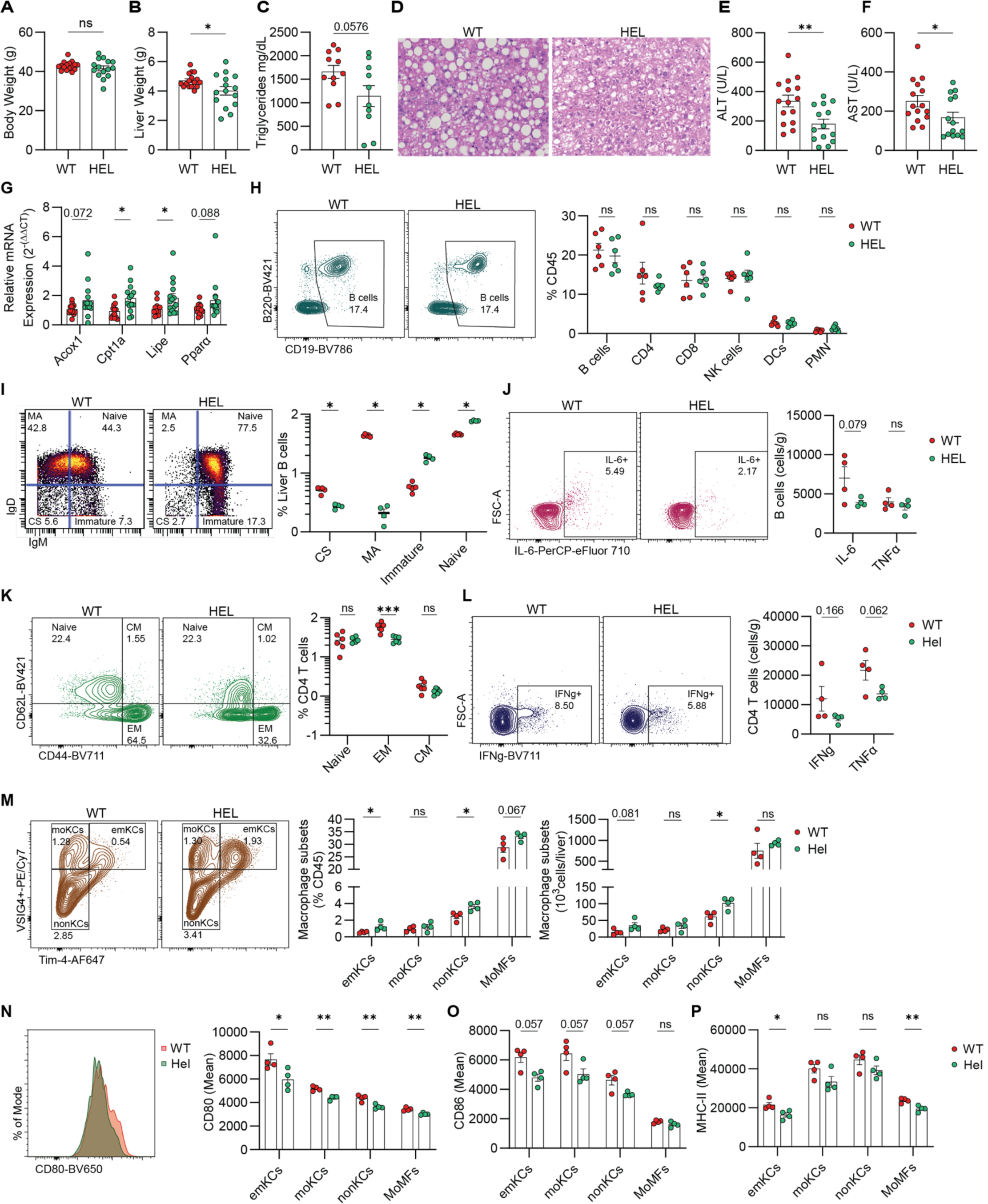
B cell receptor restriction ameliorates MASH progression. **(A-P)** WT and HEL transgenic were fed the MASH-inducing HFHC diet for 15-19 weeks. **(A)** Body weight. **(B)** Liver weight. **(C)** Liver triglyceride content. **(D)** H&E liver stain (×200 magnification). **(E)** Serum ALT levels. **(F)** Serum AST levels. **(G)** qRT-PCR gene-expression analysis of genes involved in fatty acid oxidation in the liver. **(H)** Representative flow cytometry plot showing intrahepatic B cell gating (left) and frequency of B cells, CD4 & CD8 T cells, NK cells, DCs, and PMNs (right). **(I)** Representative CyTOF plot showing class-switched (CS), mature activated (MA), naïve, and immature B cells (far left) and frequency quantification (right). **(J)** Representative flow plot of IL-6^+^ intrahepatic B cells (left) and quantification of IL-6^+^ and TNF-α^+^ intrahepatic B cells (right) after a 5-hour stimulation with PMA. **(K)** Representative flow plot of naïve, EM, and CM CD4 T cells (left) and frequency (right). **(L)** Representative flow plot of IFN-γ^+^ CD4 T cells (left) and quantification of IFN-γ^+^ and TNF-α^+^ CD4 T cells (right) post-PMA stimulation. **(M)** Representative flow cytometry plot showing macrophage subsets (left; shown populations are gated on CD45^+^ Ly6G^-^,F4/80^high^, CD11b^low/intermediate^), frequency (middle; MOMFS are identified as CD45^+^ Ly6G^-^,F4/80^low^, CD11b^high^) and number (right). **(N)** Representative MFI plot showing expression of CD80 on emKCs (left) and quantification for all macrophage subsets (right). **(O)** Mean MFI of CD86 expression by macrophage subsets. **(P)** Mean MFI of MHC-II expression by macrophage subsets. Data are representative of one independent experiment (except **A-G**, 3 independent experiments), ns = not significant, *p ≤ 0.05; **p ≤ 0.01; ***p ≤ 0.001; ****p ≤ 0.0001, unpaired t-test. Abbreviations: WT, wild-type; HEL, hen egg lysozyme; ALT, alanine transaminase; AST, aspartate aminotransferase; qRT-PCR, quantitative real-time polymerase chain reaction; NK cells, natural killer cells; DCs, dendritic cells; PMNs, polymorphonuclear neutrophils; CS, class-switched; MA, mature activated; PMA, phorbol-myristate-acetate; EM, effector memory cells, CM; central memory cells; emKCs, embryonic-derived kupffer cells; moKCs, monocyte-derived kupffer cells; nonKCs, non kupffer cells; MOMFs, monocyte-derived macrophages; MFI, mean fluorescence intensity.

### Antigen-specific B cell responses promote liver fibrosis during MASH

Previous reports from our lab indicate that intrahepatic B cells promote fibrosis during MASH^10^. However, the mechanisms through which B cells modulate fibrotic responses have been lacking. Using a CCL4 model of liver fibrosis, we observed strong interactions between HSCs and B cells via spatial transcriptomics **(Extended Data** Fig. 4A, 4B, 4C, 4D) suggesting that B cells may directly interact with HSCs to promote fibrosis. Considering that limiting BCR-mediated B cell activation during MASH protected against steatosis and inflammation, we investigated the effect of BCR restriction on liver fibrosis. HEL mice fed an HFHC diet displayed less collagen accumulation in the liver compared to controls (**Fig. 6A**). Consistently, analysis of liver tissues revealed a downregulation of the fibrosis-related genes *Col3a1* and *Tgfb1* (**Fig. 6B**). To understand how BCR restriction of B cells improves fibrosis during MASH, we assessed the antibody production by intrahepatic HEL B cells. In response to LPS, HEL B cells produced less IgG1, IgG2a, IgG2b, IgG3, and IgM, while IgA production remained unaffected (**Fig. 6C**). IgM/CD40 stimulation led to decreased IgG2a production and a surprising increase in IgA production (**Fig. 6D**), similar to that observed in the serum (**Fig. 6E**). In the liver, IgG antibodies substantially decreased (**Fig. 6F**) while IgA titers increased (**Fig. 6G**). Because HEL mice have a restricted BCR specific for HEL, we checked whether the IgA was functional by conducting bacterial flow to quantify the IgA coating of fecal bacteria. Compared to wild-type mice, HEL mice showed no difference in the IgA coating of fecal bacteria (**Fig. 6H**) indicating that HEL IgA was functional and not restricted to recognize the HEL antigen. Fc receptors for IgG have previously been reported to be expressed by HSCs^28^. Therefore, we examined the gene expression of different Fc receptors by HSCs. We found that HSCs isolated from HEL mice had significantly lower expression of *Fcmr* the gene encoding for the IgM Fc receptor and trending decreases in *Fcer1* and *Fcgr3* genes encoding for IgE and IgG Fc receptors respectively (**Fig. 6I**). Unsurprisingly, the expression of the gene encoding for the IgA Fc receptor was not different between HSCs from HEL or wild type mice (**Fig. 6I**). Together, these results suggest that antigen-specific antibody responses promote liver fibrosis during MASH through actions of B cell antibodies.

**Figure 6:**
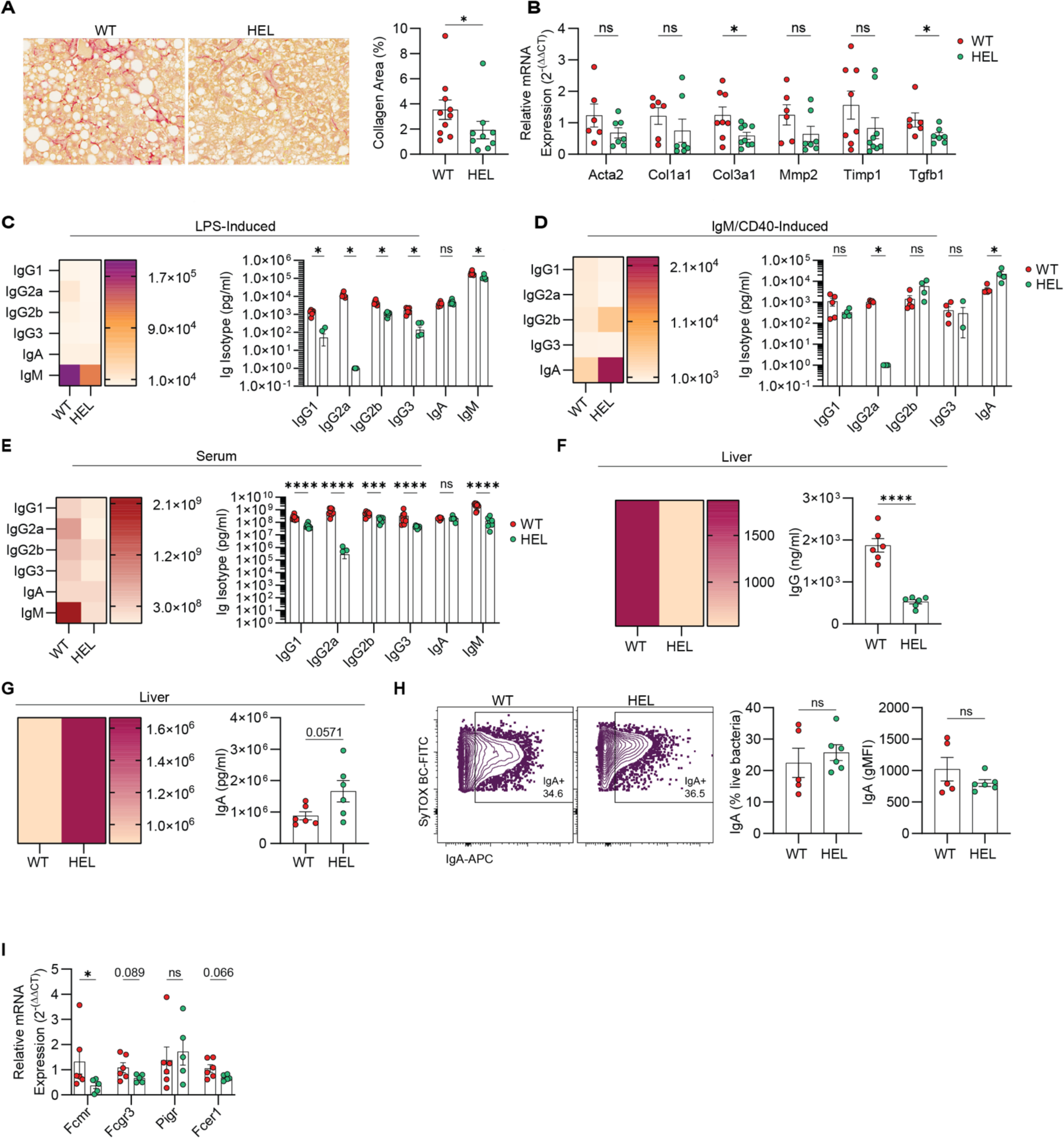
Antigen-specific B cell responses regulate liver fibrosis during MASH. **(A-I)** WT and HEL transgenic were fed the MASH-inducing HFHC diet for 15-19 weeks. **(A**) Representative picrosirius red liver stain (left; ×200 magnification) with quantification of the area with collagen deposition (right). **(B)** qRT-PCR gene-expression analysis of liver pro-fibrotic genes. **(C)** Heat map and quantification of immunoglobulin isotypes produced by HEL and WT intrahepatic B cells stimulated with LPS for 48 hours. **(D)** Heat map and quantification of immunoglobulin isotypes produced by HEL and WT intrahepatic B cells stimulated with IgM/CD40 for 48 hours. **(E)** Heat map and quantification of immunoglobulin isotypes in the sera of HEL and WT mice. **(F)** Heat map and quantification of IgG in HEL and WT liver tissue. **(G)** Heat map and quantification of IgA in HEL and WT liver tissue. **(H)** Representative flow cytometry analysis of IgA coating of fecal bacterial, (live bacterial was gated as DAPI^−^ Sytox BC^+^). **(I)** qRT-PCR gene expression analysis of Fc receptors on isolated hepatic stellate cells from HFHC-fed HEL and WT mice. Data are representative of one independent experiment, ns = not significant, *p ≤ 0.05; **p ≤ 0.01; ***p ≤ 0.001; ****p ≤ 0.0001, unpaired t-test. Abbreviations: WT, wild-type; HEL, hen egg lysozyme; LPS, lipopolysaccharide.

## DISCUSSION

B cells have been implicated in the pathogenesis of MASH, but our understanding of the mechanisms through which they promote disease is incomplete. In this report, we report that B cells accumulate in the MASH liver where they present with a more mature phenotype, corresponding to an increased abundance of class-switched B cell subsets and a naïve B cell subset characterized by an antigen inexperienced BCR. Our findings indicate the presence of immature B cells in healthy liver which diminish with MASH, suggesting the liver might serve as a site for B cell maturation. Indeed, hepatocytes have been shown to upregulate the expression of B cell activating factor^29^, which is required for B cell maturation^30^ during MASH. Additionally, several studies have reported the presence of hematopoietic stem cells that reconstitute immune cell populations in the liver under stress^31–33^, potentially differentiating locally to produce B cells^33^. Consistent with previous findings showing B cells produce a wide variety of cytokines during MASH^10, 34^, our study reveals MASH intrahepatic B cells are potent producers of polyfunctional cytokines including IL-6, Il-7, RANTES, and Granzyme B. Moreover, B cell responses are also important for MASH progression as elevated circulatory levels of IgA are associated with worse fibrosis in MASH patients^35^ while increased serum IgG antibodies against oxidative stress-derived epitopes correlate with MASH onset^29, 36^. In contrast to published reports^29^, we observed reductions in the frequency and no discernible difference in the number of antibody-producing cells in the MASH liver. Antibody production by purified intrahepatic B cells appears to be attenuated in response to TLR4 and nonspecific BCR agonism. This discrepancy may be attributed to the depletion of IgA^+^ B cells in the Peyer’s patches during MASH^37^, given that Peyer’s patches are primary sources of IgA within the liver^38^. Additionally, differences in disease stage could contribute to these findings, as disease progression has been reported to correlate with an increased presence of IgA in the liver^24^. Although local plasma cells were reduced within the MASH liver an increased deposition of IgG antibodies was observed implying increased accumulation of antibodies from extrahepatic sources. Supporting this notion, a previous study demonstrated that IgA antibodies specific for intestinal antibodies originate from the Peyer’s patches and are deposited in the liver sinusoids at steady state and during alcoholic steatohepatitis^38^.

Metabolic reprogramming of immune cells to meet energetic demand is now recognized to be crucial in regulating immune responses. Several studies have reported that activated B cells upregulate glycolysis^39^ and OXPHOS pathways to meet their energetic demand of activation^20^. Here we report that MASH intrahepatic B cells display higher ATP production rates due to increases in glycolysis and most dominantly OXPHOS. This elevated ATP production was fueled by pyruvate oxidation with minimal contribution from fatty acid oxidation. Germinal center B cells, which give rise to plasma cells have been shown to selectively utilize fatty acid oxidation to meet their metabolic demand^21^. Therefore, the observed decrease in plasma cells and local antibody production of MASH B cells may explain the lower rates of fatty acid β-oxidation seen in our model. Additionally, the preference of MASH intrahepatic B cells for using pyruvate to fuel OXPHOS may be influenced by upstream activating stimuli and the need for intermediate metabolites to launch effector responses. Indeed, nonspecific BCR stimulation of B cells led to increased glycolytic rates^39, 40^ inhibition of which led to an arrest in B cell growth^39^. Glycolysis is a major source of pyruvate within the cell, therefore an increase in glycolytic rates will lead to subsequent increases in pyruvate. Lactate, another possible source of pyruvate has been reported to be increased in the MASH liver^41^. Moreover, in activated CD4 T cells from inflamed tissue, lactate uptake is increased via SLC5A12-mediated uptake, leading to pyruvate oxidation which provides building blocks, such as citrate fatty acid synthesis, promoting cell retention in inflamed tissue^42^. Linking the metabolic rewiring of MASH B cells to its effector function, we found that blockade of pyruvate oxidation completely attenuated inflammatory cytokine responses in a manner that was dependent on BCR signaling.

In the current study, we found that restricting the BCR ameliorated MASH progression including decreased steatosis, inflammation, and fibrosis. The improvement in steatosis was attributed to enhanced fatty acid oxidation, although further research is needed to elucidate the underlying mechanisms. The amelioration of hepatic inflammation was linked to a decrease in the activation and effector responses of B cells, macrophages, and T cells. MASH is associated with substantial changes in the macrophage compartment, including loss of emKCs which are replaced by moKCs^43–45^, and the emergence of lipid-associated macrophages (LAMs)^8, 45^. Our findings suggest that B cells are involved in the reprogramming of macrophage populations during MASH. B cell-macrophage crosstalk has been reported in the liver at steady state^46^ and in the mesenteric adipose tissue during steatosis where B cells stimulate macrophages to produce pro-inflammatory cytokines in vitro^47^. In our study, BCR-restricted mice exhibited an increased abundance of emKCs and nonKCs, the population that makes up LAMs in the liver^8^. Loss of emKCs has been linked to the accumulation of moKCs, which have a more pronounced inflammatory and fibrotic phenotype^43^. Additionally, our findings suggest that B cell antigen-specific responses regulate macrophage antigen presentation to T cells which is crucial for MASH progression. Previous reports in the adipose tissue have shown that IgG antibodies stimulate macrophages to produce TNF-α by binding to their Fc-gamma receptors^48^ making the case that in addition to antigen presentation, B cell-mediated macrophage activation can promote differentiation of T cells through cytokine responses. B cell modulation of T cell activation is well established to be important for MASH progression through the promotion of hepatic inflammation via Th1 responses including the production of IFN-γ and TNF-α^10, 34^. Therefore, it is possible that BCR restriction led to a reduction in T cell-mediated hepatic inflammation as the BCR is a major upstream regulator of antigen recognition and presentation to T cells^49^.

MASH patients and experimental MASH models have elevated levels of IgG antibodies against lipid peroxidation which is associated with worsened fibrosis and disease^11, 50^. In the obese visceral adipose tissue, pathogenic IgG antibodies accumulate within the tissue and associate with fibrotic crown-like structure niches suggesting a possible role of IgG in modulating fibrosis^48^. Here, we demonstrate that restricting antigen-specific responses led to decreases in systemic and hepatic IgG levels. This reduction was accompanied by decreased expression of Fc-gamma receptors on HSCs and a subsequent decrease in liver fibrosis during MASH. Interestingly, we observed an increase in IgA in the liver of MASH mice with attenuated antigen-specific responses. A plausible explanation could be compensatory increases in polyreactive IgA antibody secretion from gut-associated lymphoid tissue in response to signaling via non-BCR routes such as TLR pathways^51, 52^. Given the pathogenic role of IgA^+^ B cells in MASH-associated HCC, it remains to be seen whether secreted antibodies play a similar immunosuppressive role^24^. In summary, we provide evidence that B cell antigen-specific responses are upregulated in MASH as evidenced by heightened local and systemic antibody responses. Notably, antigen-specific responses are sustained by pyruvate oxidation in the MASH liver, where they contribute to disease progression by promoting steatosis, hepatic inflammation, and fibrosis.

## MATERIALS AND METHODS

### Experimental Animals

Wild type (WT) C57BL/6J (Stock #000664) and HEL mice^26^ (Stock #002595) were purchased from Jackson Laboratory (Bar Harbor, ME). Mice were fed either a standard chow diet (NCD) or HFHC diet at 6 weeks of age for 15-25 weeks. The HFHC diet consisted of 40% kcal palm oil, 20% kcal fructose, and 2% cholesterol, supplemented with 42 g/L of carbohydrates in the drinking water (55% fructose, 45% sucrose; sourced from Sigma-Aldrich, St. Louis, MO)^53^. Animal experiments followed the Guide for the Care and Use of Laboratory Animals and were approved by the University of Minnesota Institutional Animal Care and Use Committee.

### MASH Assesment

Liver tissues were collected from experimental mice and 20-60mg of tissue was homogenized and quantified for triglyceride content using a colorimetric assay kit from Cayman Chemical. Serum levels of ALT and AST were assessed by the University of Minnesota’s Veterinary Medical Center Clinical Pathology Laboratory

### Immune Cell Isolation and Characterization

Intrahepatic immune cells from perfused livers were isolated through percoll gradient centrifugation as previously described^54^. Splenic immune cells were isolated via mechanical disruption on a 100µm cell strainer, followed by red blood lysis and a second filter step using a 40µm strainer to avoid clumping. Cell yield and viability were assessed with the Muse cell analyzer (CYTEK Biosciences). For seahorse and immunoassay experiments, B cells were isolated via negative selection with an EasySep B cell isolation kit (STEMCELL Technologies)

### Flow Cytometry

Immune cells were stained with zombie aqua (Biolegend; 1:200) or zombie NIR (Biolegend; 1:500) to distinguish viable from dead cells. Cells were washed with cell staining buffer (Biolegend), and non-specific binding was blocked with TruStain FcX Plus (Biolegend; 1µg). Staining for cell surface markers was performed using fluorophore-conjugated primary antibodies for 30 minutes at 4^×^C. For intracellular staining, PMA-stimulated cells were washed and stained for cell surface markers as described above. Next, a BD Cytofix/Cytoperm kit (BD Bioscience) was employed to fix and permeabilize cells before staining them with intracellular fluorophore-conjugated antibodies. Flow cytometry data were acquired on BD Fortessa X-30 H0081 (BD Biosciences) cytometer and analyzed using Flowjo software.

### CyTOF

After isolating a single cell suspension from the liver, 1-2 million single cells were stained with cisplatin (Standard BioTools; 1:2000 dilution) to discriminate viable cells. Cisplatin is washed off and non-specific binding was blocked with a 10-minute staining of cells with Fc Block (Biolegend; 1µg). Cells were stained with 0.5µg of cell surface metal-conjugated antibodies for 30 minutes at room temperature. Cells were washed and fixed with 1.6% methanol-free formaldehyde for 10 minutes at room temperature. Fixed cells were washed and resuspended in cell-ID intercalator solution (Standard BioTools; 1:4000 dilution) and kept at 4^×^C overnight. Cells are washed, counted, and prepared for acquisition at a concentration of 500,000 cells/mL. Samples were acquired on a CyTOF2 (Standard BioTools) instrument and data was analyzed using Cytobank (Beckman Coulter).

### Immunoglobulin and Cytokine Measurements

Characterization of B cell secretome was performed as follows; 1×10^6^ intrahepatic B cells were resuspended in RPMI media supplemented with 10% FBS and 1% penicillin streptomycin and primed with 5 μg/mL LPS for 16hrs in a humidified incubator set at 37^×^C, 5% CO2. After incubations, cells were washed, counted, and loaded in a mouse adaptive panel chip (IsoPlexis) and ran on the IsoLight instrument (IsoPlexis) for another 16 hours. Cytokine profiles were evaluated by IsoPlexis’s IsoSpeak software. Immunoglobulin characterizations were performed as follows: Serum samples were prepared for Legendplex assays by diluting 1:50000 for the detection of immunoglobulin isotypes using the mouse immunoglobulin isotyping kit (BioLegend). The assay was performed in a V-bottom plate and data was acquired on a BD Fortessa X-30 cytometer (BD Biosciences). Purified intrahepatic B cells were resuspended in seeded at a density of 200,000 cells per well and stimulated with 5 μg/mL LPS or 10 μg/mL IgM with 1 μg/mL CD40 with or without the addition of 2 μM UK5099 for 48hrs. The culture supernatant was collected and assessed for immunoglobulins as described above ELISA measurements were performed as follows. Purified intrahepatic B cells were stimulated with LPS, IgM/CD40, and UK5099 as described above. Supernatants were collected and assessed for TNF-α and IL-6 using deluxe ELISA kits from Biolegend. 10mg of perfused liver tissue was rinsed with cold PBS, minced, and homogenized in PBS. The homogenate was centrifuged at 5000 x g for 5 minutes at 4^×^C. The supernatant was collected and measured for IgA (Cohesion Biosciences) and IgG (Antibodies-online) via ELISA. For intracellular detection of cytokines via flow cytometry, 1×10^6^ immune cells resuspended in RPMI media supplemented with 10% FBS and 1% penicillin-streptomycin were stimulated with 2µL/mL PMA plus protein transport inhibitor (eBioscience) for 5 hours in a humidified incubator set at 37^×^C, 5% CO_2_.

### Metabolism Assays

Seahorse assays were performed on purified B cells resuspended in Seahorse RPMI media supplemented with 11 mM glucose, 1 mM pyruvate, and 2 mM glutamine. 200,000 B cells were seeded in a 96-well poly-d-lysine coated seahorse cell culture microplate. Cells were allowed to rest in a non-CO_2_ incubator at 37°C for an hour. For the Real-time ATP rate assay, measurements were performed under basal conditions and after the serial addition of 1.5 μM oligomycin and 0.5 μM rotenone/antimycin A. For Mito stress tests, 1.5 μM oligomycin, 2 μM FCCP, and 0.5 μM rotenone/antimycin A were used. For the inhibitor oxidation stress tests, we used 4 μM Etomoxir, 2 μM UK5099, 3 μM BPTES, and 500nM Thioridazine. For FAO Blue characterization of fatty acid β oxidation, 150,000 purified intrahepatic B cells were seeded on a 4-well IBIDI culture plate in a serum-free culture media containing FAOBlue detection reagent at a concentration of 5 μM. Cells were incubated for 30 minutes in a 37^×^C, 5% CO_2_ incubator. Following, cells were imaged under live conditions at 405 nm excitation.

### Quantitative Real-Time PCR

Liver tissue was harvested from mice and 20-30mg chunks were subjected to total RNA extraction using the RNeasy Plus Mini kit (Qiagen), and subsequent cDNA preparation was carried out with the iScript cDNA Synthesis kit (Bio-Rad). cDNA was quantified using SYBR Green Supermix (Bio-Rad) on a CFX Connect qPCR machine (Bio-Rad). Normalization of gene expression to GAPDH was performed, and changes in gene expression were determined using the 2(-ΔΔCT) method^55^.

### Histology

Liver chunks fixed in 10% formalin underwent histological assessment for steatosis and fibrosis through Picrosirius Red and hematoxylin and eosin (H&E) staining, conducted by the Biorepository & Laboratory Services at the UMN Clinical and Translational Science Institute. The collagen volume fraction in picrosirius red-stained images at 10X resolution was quantified using the Weka Trainable Segmentation plugin in FIJI software (ImageJ). Approximately ten different sections per histological slice were subjected to analysis. The segmentation of red staining representing collagen deposition from other colors was achieved using the plugin classifier. Subsequently, images were converted to RGB color, and the segmented color of collagen was isolated through color thresholding. The percentage area of collagen deposition was then calculated for each section using binary processing.

### Hepatic stellate cell Isolation

HSCs were isolated from healthy C57BL/6J mice as previously described^56^. Briefly, mice were anesthetized with ketamine before in situ perfusion through the portal vein. Collagenase and pronase were injected through the portal vein to digest the liver. After perfusion, in vitro, digestion was performed with collagenase/pronase solution. The liver cell suspension is filtered using a 70 μm strainer followed by a nycodenz density gradient separation. HSCs were carefully collected from the middle layer after density gradient separation. Isolated HSCs were then subjected to downstream applications.

### Statistical Analyses

Results are expressed as mean +/− SEM, and the statistical test employed for each figure is specified in the respective legend. Normality tests were conducted for all analyses using the Shapiro-Wilk test. For comparing two groups with normal distribution, p-values were determined using an unpaired, two-tailed t-test, while Mann-Whitney U tests were used for non-normal distribution. For experiments with four groups, one-way ANOVA with Fisher’s LSD test was performed to determine differences. Normal distribution (Shapiro-Wilk test) and equal variances (Brown-Forsythe) were confirmed for all datasets analyzed by ANOVA notation as follows: ns = not significant; *p ≤ 0.05; **p ≤ 0.01; ***p ≤ 0.001; ****p ≤ 0.0001. Statistical analyses were conducted using GraphPad Prism software, version (10.1.0)

## AUTHOR CONTRIBUTIONS

X.S.R., and F.B., conceived the study and designed the experiments. X.S.R. and F.B. interpreted results. F.B., H.W., and E.K., generated figures and tables. F.B. wrote the original manuscript draft and X.S.R reviewed and provided edits. F.B., H.W., G.F., K.F., E.C., S.K., P.P., and X.S.R. performed the experiments. E.R. analyzed data. X.S.R. obtained funding for, supervised, and led the overall execution of the study.

## ACKNOWLEDGMENTS

This work is supported by the National Institute of Diabetes and Digestive and Kidney Diseases (DK122056 to X.S.R.) and the National Heart, Lung, and Blood Institute (R01HL155993 to X.S.R.). F.B. is supported by an American Heart Association Predoctoral Fellowship (916054). We express our gratitude to the dedicated staff at Research Animal Resources, University Flow Cytometry Resource, Mass Cytometry Facility, Clinical and Translational Science Institute, Veterinary Medical Center Clinical Pathology Laboratory, and the Genomics Center at the University of Minnesota for their invaluable assistance.

