## Supplemental Materials for "Pyruvate Oxidation Sustains B Cell Antigen-Specific Activation to Exacerbate MASH"

### Supporting Information

#### Pyruvate Oxidation Sustains B cell Antigen-specific Responses to Exacerbate Metabolic dysfunction-associated Steatohepatitis

Fanta Barrow[1], Haiguang Wang[1], Gavin Fredrickson[1], Kira Florczak[1], Erin Ciske[1], Shalil Khanal[3], Preethy Parthiban[1], Huy Nguyen[1], Enrique Rois[1], Enis Kostallari[3], and Xavier S. Revelo[1,2,4]

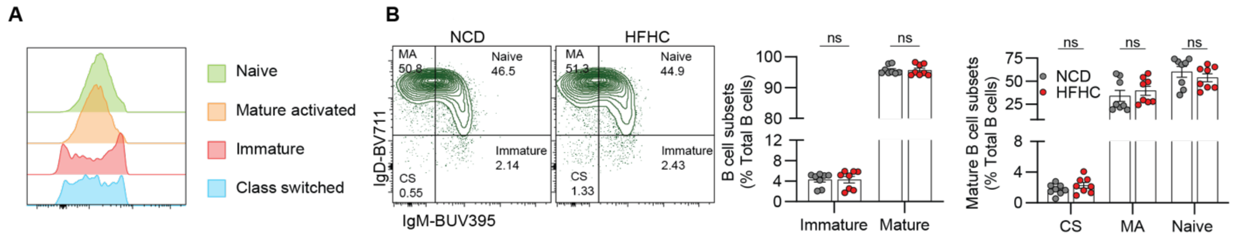


**Extended Data Fig. 1: Related to Figure 1. (A-B)** C57BL/6J mice were fed an NCD or MASH-inducing HFHC diet for 15-19 weeks. **(A)** Representative flow cytometry plot showing CD24 expression on different intrahepatic B cell subsets. **(B)** Representative flow cytometry plot showing splenic class-switched (CS), mature activated (MA), naïve, and immature B cell subsets (far left); followed by frequency of total mature (CS, MA, and naïve) and immature B cell subsets (middle left), and frequency of mature B cell subsets (right). Data are representative of one independent experiment, ns = not significant, *p ≤ 0.05; **p ≤ 0.01; ***p ≤ 0.001; ****p ≤ 0.0001, unpaired t-test. Abbreviations: NCD, normal chow diet; HFHC, high-fat high-carbohydrate.

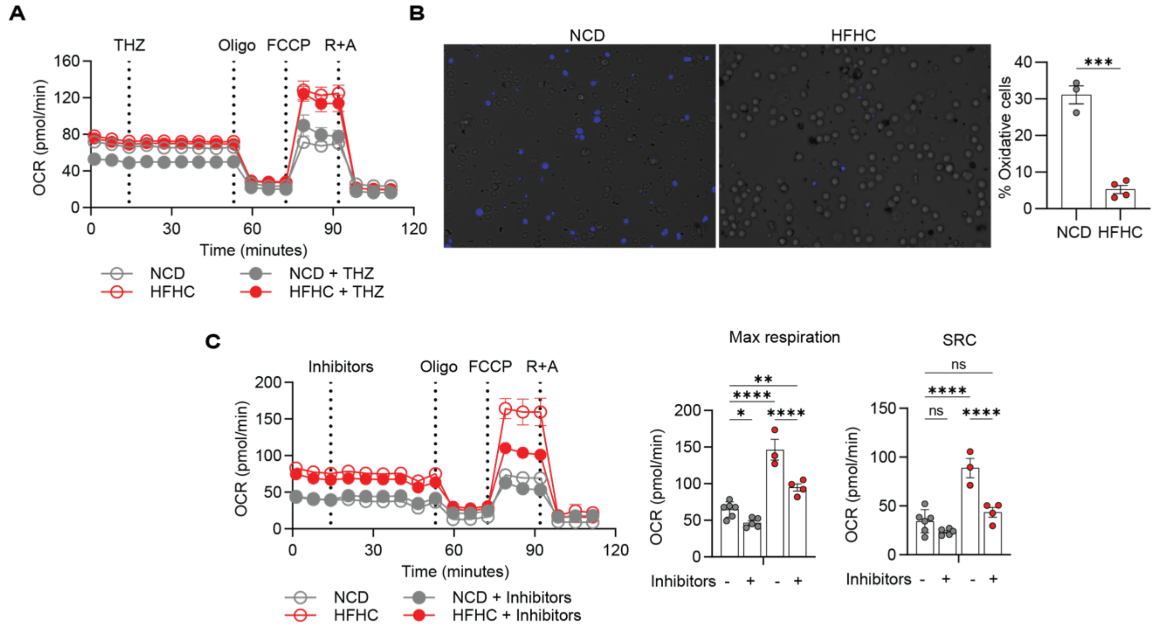


**Extended Data Fig. 2: Related to Figure 3. A-C** Data is from C57BL/6J mice fed an NCD or HFHC diet for 15-21 weeks. **(A)** Representative trace and quantification of basal, ATP-linked, maximal, and SRC of intrahepatic B cells subjected to a substrate oxidation stress test with the peroxisomal fatty acid oxidation inhibitor Thioridazine (THZ). **(B)** Representative images of Intrahepatic B cells from NCD and HFHC livers were treated with FAO Blue reagent to allow visualization of fatty acid oxidation cycles (left) and quantification of the percentage of oxidative cells per sample (left; determined by calculating the number of fluorescent cells per an average of 4 field of views per well). **(C)** Representative trace and quantification of maximal, and SRC of intrahepatic B cells subjected to a substrate oxidation stress test where three inhibitors (Etomoxir, BPTES, and UK5099) are tested. Results represent the mean of one independent experiment. Bars represent mean +/− SEM; ns = not significant; *p ≤ 0.05; **p ≤ 0.01; ***p ≤ 0.001; ****p ≤ 0.0001 by One-Way ANOVA or an unpaired t-test. Abbreviations: NCD, normal chow diet; HFHC, high-fat high-carbohydrate; Oligo, oligomycin; R+A, rotenone, and antimycin A; SRC, spare respiratory capacity; FCCP, Carbonyl cyanide-4 (trifluoromethoxy) phenylhydrazone; SRC, spare respiratory capacity.

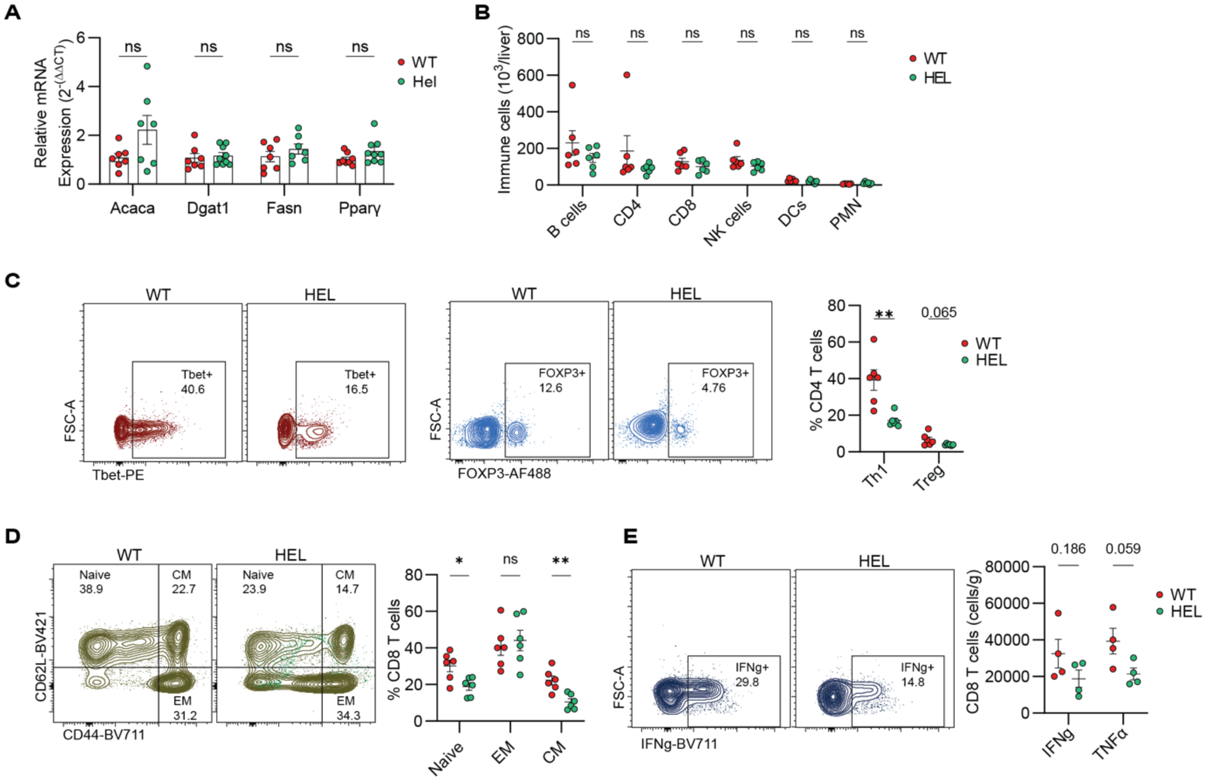


**Extended Data Fig. 3: Related to Figure 5. (A-E)** WT and HEL transgenic were fed the MASH-inducing HFHC diet for 15-19 weeks. **(A)** qRT-PCR gene-expression analysis of genes involved in lipogenesis in the liver. **(B)** Number of B cells, CD4 & CD8 T cells, NK cells, DCs, and PMNs in WT and HEL livers. **(C)** Representative flow plot of $Tbet^{+}$ Th1 cells (left), $FOXP3^{+}$ Tregs (middle), and frequency (right) in the liver. **(D)** Representative flow plot of naïve, EM, and CM CD8 T cells (left) and frequency (right). **(E)** Representative flow plot of IFN-γ+ CD8 T cells (left) and quantification of IFN-γ+ and TNF-α+ CD8 T cells (right) post-PMA stimulation. Data are representative of one independent experiment, ns = not significant, *$p \leq 0.05$; **$p \leq 0.01$; ***$p \leq 0.001$; ****$p \leq 0.0001$, unpaired t-test. Abbreviations: WT, wild-type; HEL, hen egg lysozyme; PMA, phorbol-myristate-acetate; EM, effector memory cells, CM; central memory cells.

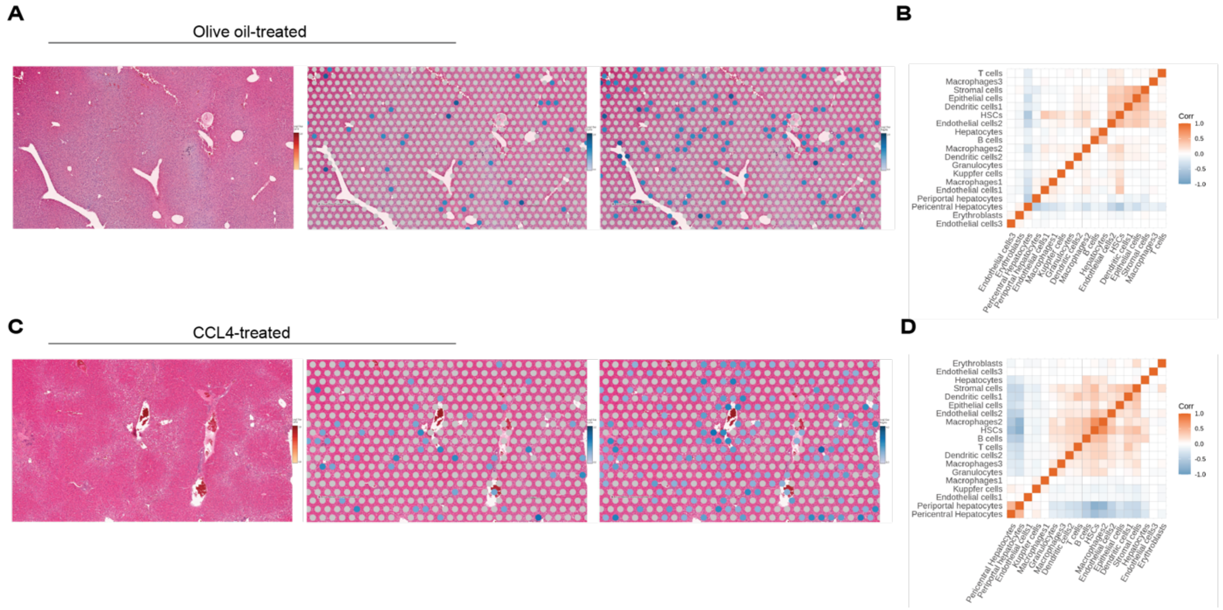


**Extended Data Fig. 4: Related to Figure 6. (A-D)** Spatial transcriptomic data from CCL4-induced fibrosis mouse model. **(A)** H&E stains of olive oil-treated liver sections showing CD19 staining (left), B cell localization mapping (middle), and HSC localization mapping (right). **(B)** Neighborhood analysis of cell-cell interactions in olive oil-treated livers. **(C)** H&E stains of CCL4-treated liver sections showing CD19 staining (left), B cell localization mapping (middle), and HSC localization mapping (right). **(D)** Neighborhood analysis of cell-cell interactions in CCL4-treated livers. Data are representative of one independent experiment. Abbreviations: H&E, hematoxylin, and eosin; HSC, hepatic stellate cell.

**Extended Data Table 1: Fluorophore-Conjugated Antibodies**

| Marker | Fluorophore | Clone | Supplier |
|---|---|---|---|
| CD45.2 | BV510 | 104 | Biolegend |
| CD3 | BUV737 | 145-2C11 | BD Bioscience |
| CD4 | BUV395 | RM4-5 | BD Bioscience |
| CD8a | BUV805 | 53-6.7 | BD Bioscience |
| CD11b | PE/Cy7 | M1/70 | Biolegend |
| CD19 | BV785 | 6D5 | Biolegend |
| B220 | BV421 | RA3-6B2 | Biolegend |
| IgM | BUV395 | R6-60.2 | BD Bioscience |
| IgD | BV711 | 11-26c.2a | Biolegend |
| IgD | PerCP/Cy5.5 | 11-26c.2a | Biolegend |
| CD24 | APC-efluor780 | M1/69 | Invitrogen |
| CD138 | PE/Dazzle 594 | 281-2 | Biolegend |
| IgA | APC | mA-6E1 | Invitrogen |
| NK1.1 | APC | PK136 | Biolegend |
| CD11c | BV711 | N418 | Biolegend |
| Ly6G | PE | 1A8 | Biolegend |
| F4/80 | FITC | BM8 | Biolegend |
| CD44 | BV711 | IM7 | Biolegend |
| CD62L | BV605 | GL-3 | Biolegend |
| CD86 | PE | GL-1 | Biolegend |
| CD80 | BV650 | 16-10A1 | Biolegend |
| MHC II | FITC | MS/114.15.2 | Biolegend |
| CD40 | PerCP-Cy5.5 | 3/23 | Biolegend |
| IL-6 | PerCP-eFluor 710 | MP5-20F3 | ThermoFisher |
| IFNγ | BV711 | XMG1.2 | Biolegend |
| TNFα | BV650 | MP6-XT22 | Biolegend |

**Extended Data Table 2: Metal-Conjugated Antibodies**

| Metals | Antibodies | Clone |
|---|---|---|
| 141Pr | Ly6G | 1A8 |
| 142Nd | CD11c | HL3 |
| 143Nd | CCR2 | 475301R |
| 144Nd | MHC Class I | 28-14-8 |
| 145Nd | CD4 | RM4-5 |
| 146Nd | CD5 | 53-7.3 |
| 147Sm | CD45.2 | 104 |
| 148Nd | CD11b | M1/70 |
| 149Sm | CD19 | 6D5 |
| 150Nd | Ly-6C | HK1.4 |
| 151Eu | IgM | RMM-1 |

| | | |
|---|---|---|
| 152Sm | CD3e | 145-2C11 |
| 153Eu | CD64 | X54-5/7.1 |
| 154Sm | Tim-4 | RMT4-54 |
| 156Gd | IgD | 11-26c.2a |
| 159Tb | F4/80 | BM8 |
| 160Gd | CD62L | MEL-14 |
| 168Er | CD8a | 53-6.7 |
| 169Tm | CD44 | IM7 |
| 170Er | NK1.1 | PK136 |
| 172Yb | CD86 | GL1 |
| 173Yb | CD138 | 281-2 |
| 174Yb | I-A/I-E | MS/114.15.2 |
| 176Yb | CD45R (B220) | RA3-6B2 |

**Extended Data Table 3: qRT-PCR Primers**

| Primer | Sequence |
|---|---|
| *B-actin* | Forward: 5′-TGTTACCAACTGGGACGACA-3′<br>Reverse: 5′-CTTTTCACGGTTGGCCTTAG-3′ |
| *Gapdh* | Forward: 5′-AACGACCCCTTCATTGAC-3′<br>Reverse: 5′-TCCACGACATACTCAGCAC-3′ |
| *Acox1* | Forward: 5’-GTCTCCGTCATGAATCCCGA-3’<br>Reverse: 5’-TGCGATGCCAAATTCCCTGA-3’ |
| *Cpt1a* | Forward: 5’-GGTCTTCTCGGGTCGAAAGC-3’<br>Reverse: 5’-TCCTCCCACCAGTCACTCAC-3’ |
| *Lipe* | Forward: 5’-ACGCTACACAAAGGCTGCTT-3’<br>Reverse: 5’-TCGTTGCGTTTGTAGTGCTC-3’ |
| *Ppara* | Forward: 5’-TATTCGGCTGAAGCTGGTGTAC-3’<br>Reverse: 5’-CTGGCATTTGTTCCGGTTCT-3’ |
| *Acaca* | Forward: 5’-CGCTCAGGTCACCAAAAAGAAT-3’<br>Reverse: 5’-GTCCCGGCCACATAACTGAT-3’ |
| *Dgat1* | Forward: 5’-GGAATATCCCCGTGCACAA-3’<br>Reverse: 5’-CATTTGCTGCTGCCATGTC-3’ |
| *Fasn* | Forward: 5’-TCCTGGAACGAGAACACGATCT-3’<br>Reverse: 5’-GAGACGTGTCACTCCTGGACTTG-3’ |
| *Pparg* | Forward: 5’-TCGCTGATGCACTGCCTATG-3’<br>Reverse: 5’-GAGAGGTCCACAGAGCTGATT-3’ |
| *Acta2* | Forward: 5′-GGCTCTGGGCTCTGTAAGG-3′<br>Reverse: 5′-CTCTTGCTCTGGGCTTCATC-3′ |
| *Col1a1* | Forward: 5′-ACATGTTCAGCTTTGTGGACC-3′<br>Reverse: 5′-TAGGCCATTGTGTATGCAGC-3′ |
| *Col3a1* | Forward: 5’-CTGTAACATGGAAACTGGGGAAA-3’<br>Reverse: 5’-CCATAGCTGAACTGAAAACCACC-3’ |
| *Mmp2* | Forward: 5′-TTCCCCCGCAAGCCCAAGTG-3′<br>Reverse: 5′-GAGAAAAGCGCAGCGGAGTGACG-3′ |
| *Timp1* | Forward: 5′-GCATCTCTGGCATCTGGCATC-3′<br>Reverse: 5′-GCGGTTCTGGGACTTGTGGGC-3′ |
| *Tgfb1* | Forward: 5′-GGTTCATGTCATGGATGGTGC-3′<br>Reverse: 5′-TGACGTCACTGGAGTTGTACGG-3′ |
